# Stereochemistry-Aware Drug-Target Affinity Prediction

**DOI:** 10.64898/2026.05.14.725200

**Authors:** Santiago Ferreyra, Iago Dutra, Aldo Galeano, Alberto Paccanaro

## Abstract

Drug–target affinity (DTA) prediction is a key task in drug discovery, enabling the estimation of the interaction strength between candidate compounds and biological targets. However, current models rely on connectivity-based molecular representations and do not explicitly account for the spatial organization, also known as stereochemistry. This limitation becomes evident when considering chirality, where a drug can exist as enantiomers, i.e., molecules that share the same atoms and bonds but differ in their three-dimensional arrangement. Despite their chemical similarity, they can interact differently with the same target, leading to variations in binding affinity and biological activity. In this paper, we propose a stereochemistry-aware DTA prediction framework that incorporates this information into molecular representations. Drug representations are learned from chemical structure using a directed-bond message passing graph neural network that captures enantiomers configurations, while protein targets are represented through sequence-based embeddings. Experiments on the Davis dataset demonstrate that our model can improve affinity prediction. Importantly, a case study on a manually curated dataset of enantiomers with different biological action shows that the model is able to distinguish the affinities in the two forms consistent with their experimentally observed biological activity. These findings support the relevance of stereochemistry-aware molecular representation for more accurate and chemically faithful DTA prediction.

## 1 Introduction

As drug discovery continues to evolve, increasing attention has been devoted to computational approaches that support tasks such as drug-drug interaction analysis, drug repurposing, and synergy prediction (Kairys et al., 2019). Within this landscape, drug-target affinity (DTA) prediction plays a central role, as it aims to quantify the binding strength between a compound and its target protein, providing a more informative signal than binary interaction labels (Oprea and Mestres, 2012). Accurate estimation of this affinity is essential for prioritizing candidate molecules, guiding lead optimization, and informing downstream pharmacological decisions (Spassov, 2024).

Despite its importance, experimental determination of binding affinity through techniques such as high-throughput screening (HTS) or biophysical assays remains costly and time-consuming, limiting scalability (Kairys et al., 2019; Thafar et al., 2019). Classical computational approaches, including molecular docking (Trott and Olson, 2010) or molecular dynamics (MD) simulations (Liu et al., 2018), offer viable alternatives but often entail substantial computational overhead. Consequently, deep learning models have gained prominence for enabling efficient and scalable affinity prediction in large chemical and proteomic spaces.

Recent advancements have led to a variety of deep learning-based methods for DTA prediction, such as drug-target association networks, convolutional neural networks (CNNs) (Shim et al., 2021), graph neural networks (GNNs) (Shah et al., 2025), and transformer-based models (Liu et al., 2018). In general, these methods aim to learn informative representations of both molecular compounds and protein targets aiming to capture relational patterns that govern binding interactions.

Nevertheless, molecular representation remains a critical limitation in many existing models. In particular, most approaches encode compounds primarily from molecular topology, using 2D molecular graphs or SMILES-based encodings (Öztürk et al., 2018; Nguyen et al., 2021; Shah et al., 2025). Although these representations capture which atoms are connected, they may not fully capture information on how atoms are arranged in three-dimensional space. This is biologically relevant because molecular recognition is inherently three-dimensional, as a ligand must fit into a protein binding pocket with a compatible spatial arrangement of atoms and functional groups.

This distinction is especially important for enantiomers, molecules that share the same molecular formula and atomic connectivity but differ in spatial configuration (Somogyi et al., 2004). When a compound occurs in two stereochemical forms, they can exhibit different interactions despite appearing equivalent from a connectivity-based perspective. A representative example is ibuprofen, which exists as two enantiomeric forms, S-ibuprofen and R-ibuprofen, illustrated in Figure 1. Despite sharing the same atoms and bonds, their opposite three-dimensional arrangements produce distinct interactions within the cyclooxygenase enzymes. These differences translate into different pharmacological properties: S-ibuprofen is responsible for the direct COX-inhibitory effect observed at clinically relevant concentrations, whereas R-ibuprofen shows weaker direct activity against COX enzymes (Evans, 2001).

**Figure 1.**
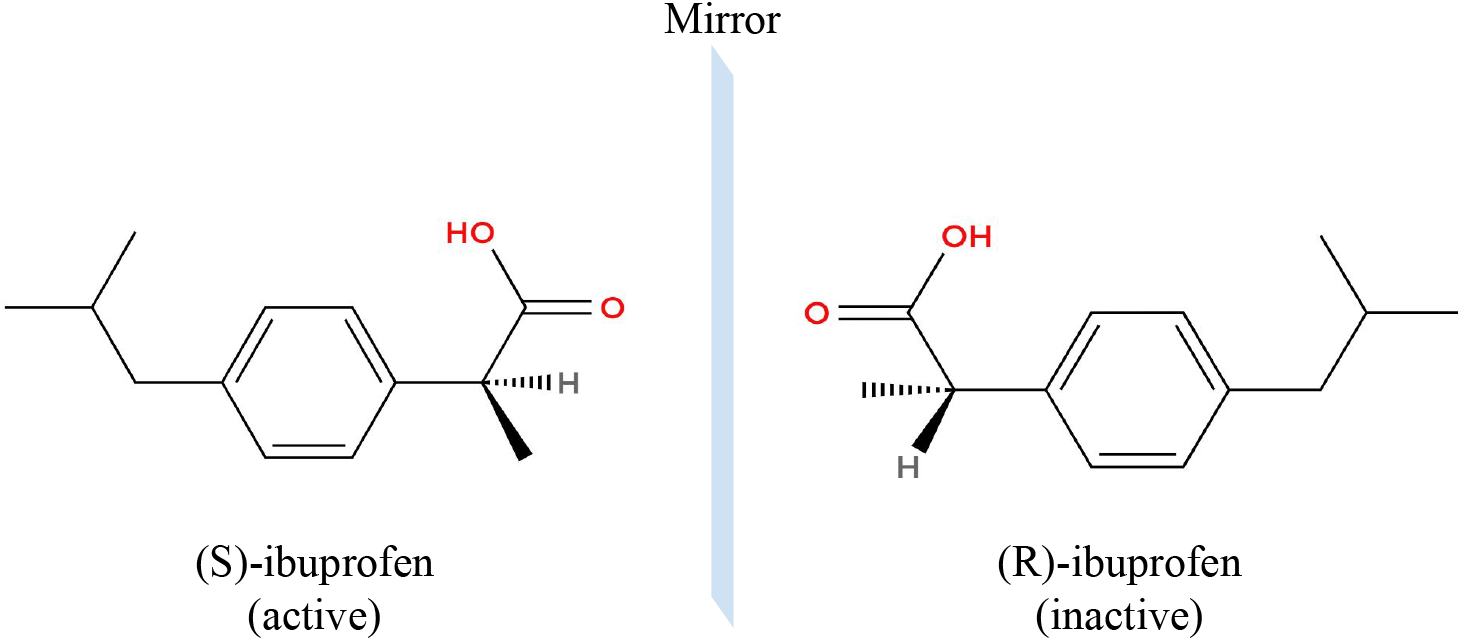
Stereochemistry. Representation of the two enantiomeric forms of ibuprofen. Solid wedges indicate bonds projecting toward the viewer, whereas dashed wedges indicate bonds projecting away from the viewer. The R- and S-forms are non-superimposable mirror images of one another and exhibit different pharmacological properties, with S-ibuprofen being the pharmacologically active form for the cyclooxygenase enzymes.

Even though this physical configuration has a high biological impact, many computational methods used in DTA prediction do not explicitly encode this stereochemical information. When molecules are represented, pairs of enantiomers may be mapped to identical or highly similar representations, limiting the ability of models to distinguish compounds with different three-dimensional configurations. As a result, models that ignore stereochemistry may produce less accurate or chemically inconsistent affinity predictions.

To address this limitation, we propose a stereochemistry-aware framework for drug–target affinity prediction. On the molecular side, compounds are encoded using a directed-bond message passing neural network (D-MPNN) that incorporates stereochemical information into the learned drug representation, allowing the model to distinguish enantiomers. On the protein side, target sequences are represented using pretrained ESM-C (ESM Team, 2024) protein language model embeddings, which provide contextual sequence-level information relevant to molecular recognition. The resulting drug and protein representations are combined to predict binding affinity, enabling the model to account for both ligand stereochemistry and target-specific biological context.

The main contributions of this paper center on the development of a stereochemistry-aware framework for drug-target affinity prediction that incorporates stereochemical molecular information into the drug representation. This approach combines directed-bond message passing for drug encoding with pretrained protein sequence embeddings for target representation. We evaluate the proposed framework on the Davis benchmark (Davis et al., 2011) and analyze stereochemical compounds, demonstrating that stereochemical information improves affinity prediction. Moreover, we present a case study on a manually curated dataset showing that our model can distinguish molecules with identical connectivity but different three-dimensional arrangements.

### 2 Methodology

### 2.1 Model Architecture

The proposed framework comprises two main components: a stereochemistry-aware molecular encoder and a protein representation module based on precomputed embeddings from ESM-C. The encoded molecular and protein features are subsequently integrated through an attention-based module, enabling joint representation learning for affinity prediction, as shown in Figure 2.

**Figure 2.**
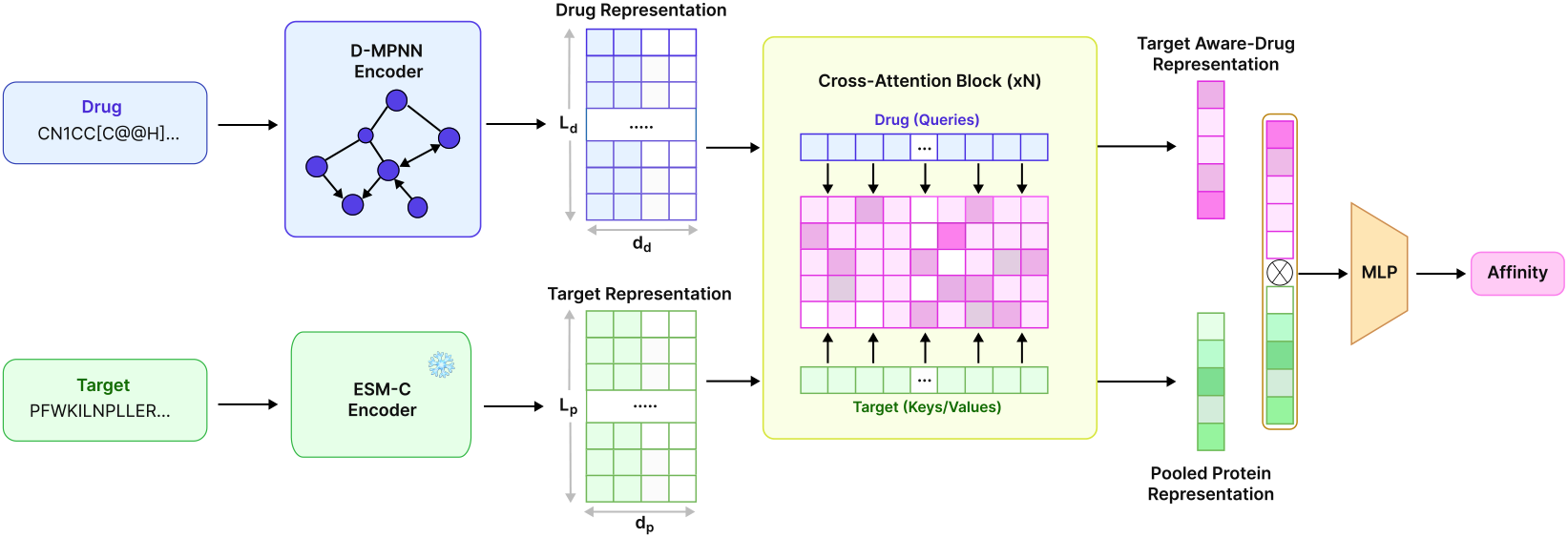
Overview of the proposed framework. Drug molecules are encoded using a directed-bond message passing neural network (D-MPNN), which operates on molecular graph information and incorporates stereochemical configuration into the learned drug representation. Target protein sequences are encoded using frozen ESM-C embeddings, providing residue-level embeddings. Drug features are used as queries, while protein features are used as keys and values in the cross-attention blocks, enabling the model to learn target-aware molecular representations. The attended drug and protein representations are pooled into fixed-size vectors and passed through a MLP to predict the binding affinity.

### 2.2 Molecular Representation

For each drug, we construct a molecular graph from its isomeric SMILES representation using RDKit (Landrum et al., 2013). Atoms are represented as nodes and covalent bonds as edges. To preserve stereochemical information, the graph is built from isomeric SMILES obtained from PubChem (Wang et al., 2009), allowing chiral and bond-configuration annotations to be reflected in the atom and bond features. Each atom *i* is associated with an initial feature vector 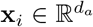, and each bond (*i, j*) is associated with a bond feature vector 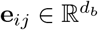.

We encode the molecular graph using a directed message passing neural network (D-MPNN). Instead of passing messages directly between atoms, our model performs message passing over directed bonds. For each undirected bond (*i, j*), two directed edges *i* →*j* and *j*→ *i* are introduced. This construction makes the message associated with a bond direction-dependent: information propagated from *i* to *j* is represented separately from information propagated from *j* to *i*. By avoiding a symmetric treatment of bonded atom pairs, the encoder can retain orientation-sensitive local contexts, which is important for representing stereochemical configurations.

The initial hidden representation of each directed bond is computed from the source atom features and the corresponding bond features:

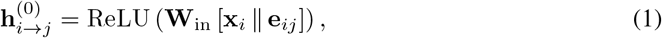

where ∥ denotes concatenation and **W**_in_ is a learnable linear transformation. In this way, each directed edge embedding encodes both atomic identity and bond-specific information from the outset.

Message passing is then performed over the directed bond graph for *T* iterations. A directed bond *i* →*j* receives information from other directed bonds that terminate at its source atom *i*. We therefore define its neighborhood as

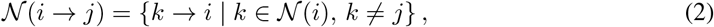

where 𝒩 (*i*) denotes the set of atoms bonded to *i*. Excluding *j* prevents immediate backtracking along the reverse edge. At iteration *t*, incoming messages are aggregated as

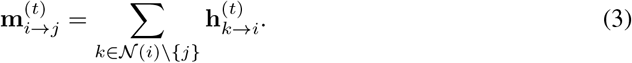

This aggregation step allows each directed bond to incorporate contextual information from its local chemical environment. The representation is then updated by combining the initial embedding with the aggregated message:

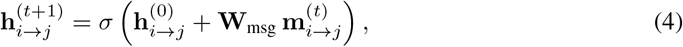

where **W**_msg_ is a learnable weight matrix shared across iterations. By repeatedly applying this update, the model progressively expands the receptive field of each directed bond representation, allowing it to encode higher-order structural dependencies.

After *T* message passing steps, we recover atom-level representations by aggregating the final directed bond embeddings that point to each atom. Specifically, we define:

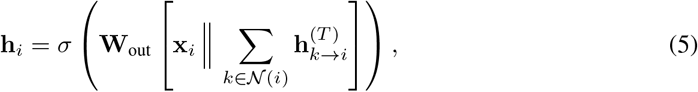

where **W**_out_ is a learnable parameter matrix. This step fuses the original atomic features with the structural information accumulated through the incident directed bonds. This is repeated by *L* iterations to get final atom-level embeddings.

### 2.3 Target Representation

On the protein side, a pretrained protein language model is employed to encode protein sequences. Each target protein is represented by its amino acid sequence, which is processed using ESM-C (600M) (ESM Team, 2024). Residue-level embeddings are extracted for each position in the sequence, yielding a representation of dimension *L*_*p*_ × *d*_*p*_, where *L*_*p*_ is the sequence length and *d*_*p*_ is the ESM-C embedding dimension.

Given the maximum input length constraint of 2048 tokens, protein sequences longer than this limit are divided into non-overlapping contiguous chunks. Each chunk is independently processed by ESM-C to generate embeddings, and the resulting representations are concatenated to reconstruct the full residue-level protein representation. This procedure ensures scalability to long sequences while preserving residue-level resolution across the entire protein, producing contextualized representations that capture evolutionary signals and functional patterns.

### 2.4 Cross-Attention Module

After obtaining atom-level drug embeddings and residue-level protein embeddings, both representations are projected into a shared latent space. The interaction between the two modalities is then modeled using a cross-attention module, where the drug representation is used as query matrix *Q*_*d*_, while protein representation is used as key and value matrices *K*_*p*_ and *V*_*p*_, respectively. The cross-attention operation is computed using scaled dot-product attention:

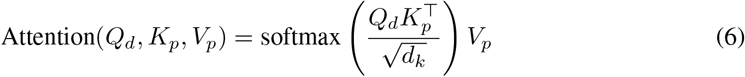

Thus, the protein representation provides target-specific context, while the output of the module is a target-aware drug representation. Protein padding masks are applied during attention so that padded residue positions do not contribute to the interaction modeling. Subsequently, the representation is aggregated by pooling over the drug atoms rather than the protein residues.

Because protein sequences are substantially longer than small-molecule graphs, the vast majority of residues lie outside the active binding pocket. Directly pooling the protein sequence would therefore dilute the localized binding signal with non-interacting background noise. By using the drug atoms to query the target sequence, the cross-attention mechanism effectively isolates the relevant interaction context, yielding a concentrated affinity representation upon drug-level readout.

### 2.5 Prediction Module

Finally, the target-aware drug representation is aggregated into a fixed-size vector using masked mean pooling over the drug atoms. In parallel, the protein representation is summarized using residue-level pooling. The resulting drug and protein are concatenated to form a joint drug-target interaction representation. This interaction vector is fed into a MLP block, which serves as a regression head to predict the drug-target binding affinity. The detailed architecture of the block is provided in Appendix.

## 3 Materials

### 3.1 Datasets

We conduct experiments on the Davis dataset, a widely adopted benchmark for drug-target affinity (DTA) prediction (Davis et al., 2011). This dataset comprises 68 small-molecule kinase inhibitors and 442 protein kinases, resulting in 30,056 drug-target interaction pairs with experimentally measured binding affinities reported as dissociation constants (*K*_*d*_). The affinity values span several orders of magnitude, which can hinder stable optimization and bias the affinity prediction. To mitigate this issue, we transform the affinity values into the logarithmic scale (*pK*_*d*_), following standard practice in the literature, as defined in Eq. (7):

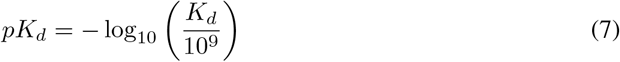

It is important to highlight that small changes in *pK*_*d*_ account for significant changes in binding affinity, given the scale of the transformation. For evaluation, we adopt the standard data splitting protocol established by GraphDTA (Nguyen et al., 2021). The dataset consists of six folds, where one fold is held out as the test set and the remaining five folds are used for training. This setup ensures a strict separation between training and testing samples, preventing information leakage and enabling a fair comparison with prior work.

Notably, 23 of the drugs, accounting for 10,174 interactions, possess explicit stereochemical annotations, though their corresponding enantiomers are not always present in the dataset. These annotations capture the three-dimensional spatial arrangements that directly influence molecular conformation and binding affinity. More details are provided in the Appendix.

Finally, to evaluate the model’s capacity to differentiate between such drugs, we curated a dataset comprising pharmacologically relevant chiral compounds sourced from PubChem (Wang et al., 2009), alongside their respective target proteins retrieved from UniProt (The UniProt Consortium, 2023).

### 3.2 Metrics

To evaluate the performance of the proposed model, we employ four metrics: Mean Squared Error (MSE), Concordance Index (CI), Mean Reversion Coefficient 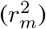, and Area Under the Precision-Recall Curve (AUPR). These metrics capture both regression accuracy and ranking consistency, as well as the model’s ability to distinguish high-affinity interactions.

- **Mean Squared Error (MSE)**. Quantifies the prediction error by measuring the average squared difference between predicted and true affinity values:

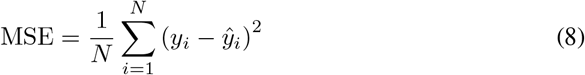
- **The Concordance Index (CI)**. Measures the consistency of ranking between predicted and true affinities of correctly ordered pairs among all comparable pairs (*i, j*) such that *y*_*i*_ *> y*_*j*_:

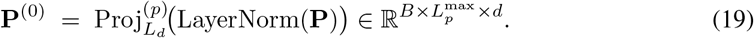

where *Z* denotes the total number of comparable pairs, and the indicator function *h*(*x*) is defined as:

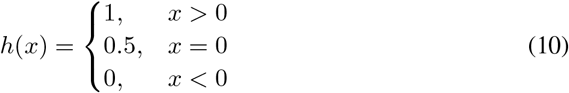
- **Mean Reversion Coefficient** 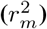. Evaluates external predictive reliability of the model and penalizes deviations from the ideal regression line:

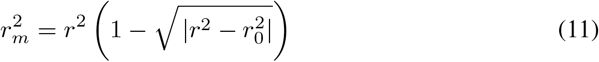

where *r*^2^ is the squared correlation coefficient between observed and predicted values, and 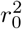 is the squared correlation coefficient for the regression through the origin.
- **Area Under the Precision-Recall Curve (AUPR)**. Summarizes the trade-off between precision and recall across different decision thresholds. To calculate this, we binarize the *pK*_*d*_ values using 7 as a threshold, following common practice (Shah et al., 2025).

## 4 Results

In this section, we evaluate the proposed framework on the Davis dataset and compare its performance against representative baseline methods. The model is trained using the Mean Squared Error (MSE) loss between predicted and experimental affinity values. In addition to the standard benchmark evaluation, we conduct case studies on enantiomeric compound pairs to assess whether the proposed model and baseline methods can distinguish affinity differences in stereochemically sensitive scenarios.

### 4.1 Baselines

We compare our proposed model with five representative baseline methods, covering traditional approaches, early deep learning models, and the latest multi-scale frameworks:

- **GraphDTA (Nguyen et al., 2021)**: leveraged the structural features of drugs meaning molecular graphs and one-hot encoding of protein sequences. A 3-layer GNN (GCN, GAT, GIN, GAT-GCN) for the extraction of structural features for drugs, a 3-layer 1D-CNN for protein sequences and FC as head was used to predict DTA.
- **AttentionDTA (Zhao et al., 2022)**: utilizes a label encoder, embedding layer and a 1D-CNN block to extract features from both drugs and proteins separately. A bilateral multi-head attention and a multi layer perceptron layer is used for the head of the framework to predict the affinity.
- **GDilatedDTA (Zhang et al., 2024)**: introduces a graph dilated convolution strategy to improve the representation learning of drug-target pairs. The model first encodes drugs and target sequences, and then applies dilated convolutional operations to capture broader contextual information and latent interaction patterns for DTA prediction.
- **GEFormerDTA (Liu et al., 2024)**: proposes a framework that considers bond encoding, degree centrality encoding, spatial encoding of drug molecule graphs, and the structural information of proteins such as secondary structure and accessible surface area. This method uses a graph early fusion module that fuses drugs and protein graphs and applies GCN to obtain graph-based representation of proteins. Information in this framework is passed between protein and drug pipelines to learn the interaction better.
- **DeepDTAGen (Shah et al., 2025)**: multitask deep learning framework designed to predict drug-target affinity and generate target-aware drug molecules. In its DTA prediction branch, compounds are represented using SMILES and molecular graph features, which are encoded through graph convolutional network (GCN) layers. Target proteins are represented using word embeddings followed by gated CNN layers to capture sequence-level features. The resulting drug and protein representations are integrated and passed to a fully connected module to predict the continuous affinity values.

### 4.2 Experiments

We evaluate our model against representative baselines on the Davis dataset. As shown in Table 1, our method obtains the best performance in terms of MSE, achieving 0.204 ±0.009, which indicates lower regression error than all compared baselines. Our method also achieves the highest reported AUPR, with 0.781 ±0.020, suggesting improved ranking performance for identifying high-affinity drug-target pairs. For the remaining metrics, our method obtains the second-best performance: a CI of 0.894 ± 0.005, only slightly below GEFormerDTA, and an 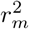 of 0.695 ± 0.019, below DeepDTAGen but higher than the other reported baselines. Overall, these results show that the proposed architecture improves affinity regression accuracy and high-affinity ranking while maintaining strong ranking performance.

**Table 1:** Performance comparison on the Davis dataset. The best results are in **bold** and second-best results are <u>underlined</u>. We take the values for baseline models from their respective original papers. Absent values represent metrics not reported by the original authors.

| Model | ↓ MSE (Std) | ↑ CI (Std) | ↑ $r_m^2$ (Std) | ↑ AUPR (Std) |
| --- | --- | --- | --- | --- |
| GraphDTA | 0.229 | 0.893 | 0.663 | — |
| AttentionDTA | $0.216 \pm 0.019$ | $0.887 \pm 0.005$ | $0.677 \pm 0.024$ | <u>$0.776 \pm 0.024$</u> |
| GDilatedDTA | 0.237 | 0.885 | 0.686 | — |
| GEFormerDTA | <u>0.212</u> | **0.895** | — | — |
| DeepDTAGen | $0.214 \pm 0.006$ | $0.890 \pm 0.004$ | **$0.705 \pm 0.001$** | $0.772 \pm 0.002$ |
| **OurMethod** | **$0.204 \pm 0.009$** | <u>$0.894 \pm 0.005$</u> | <u>$0.695 \pm 0.019$</u> | **$0.781 \pm 0.020$** |

Figure 3 complements the quantitative evaluation by illustrating the joint density distribution of experimental versus predicted affinities. The alignment of density contours along the diagonal demonstrates that the model successfully captures overall binding trends within the Davis test set, while off-diagonal dispersion reflects residual prediction errors. Importantly, the Davis benchmark consists exclusively of kinase targets, with *pK*_*d*_ values bounded between 5 and 10.8 and disproportionately concentrated near 5. This restricted dynamic range and narrow target scope inherently limit the generalizability of models trained on this dataset to broader pharmacological contexts.

**Figure 3.**
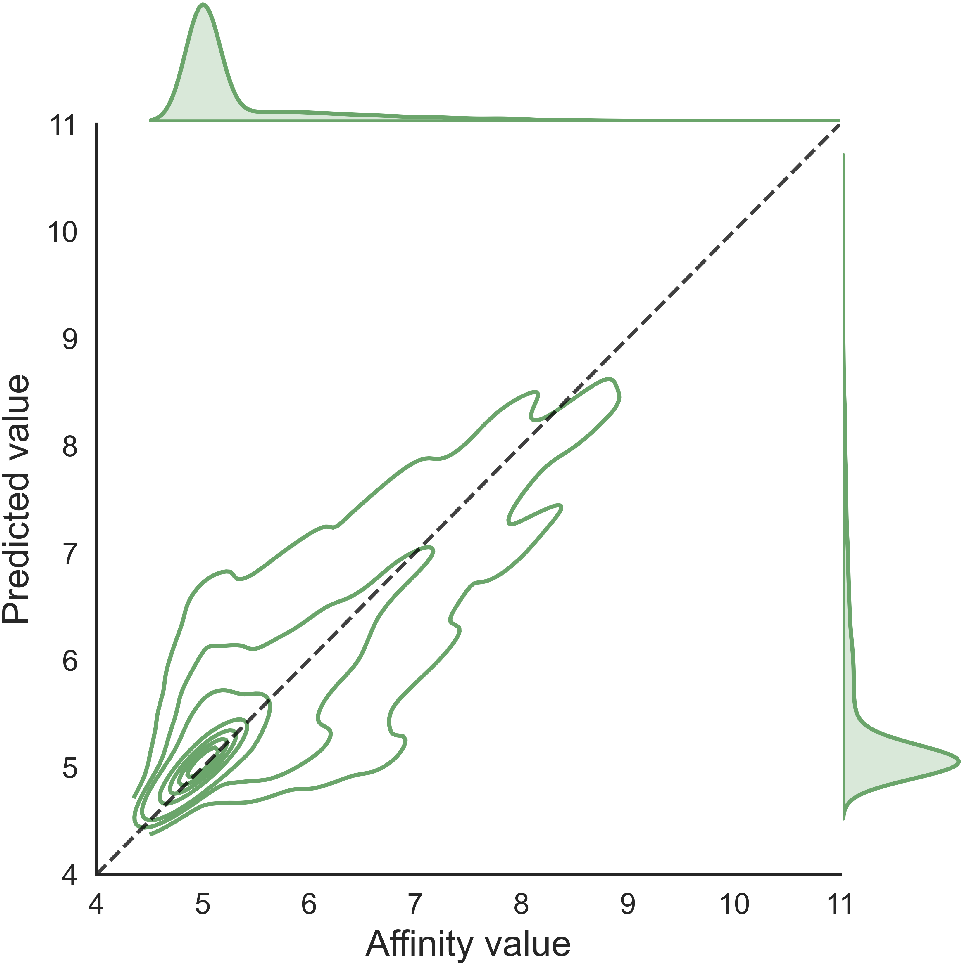
Density-level analysis of predicted and experimental affinities on the Davis test set. The contour curves represent kernel density estimation levels over the joint distribution of experimental affinity values and model-predicted affinity values.

## 5 Case Study

To assess whether the model can discriminate between stereoisomeric forms, we evaluated it on a manually curated set of 10 pharmacologically relevant isomeric drug pairs that are not present in the Davis dataset. Because experimentally validated affinity measurements for both isomers against the same protein target are not consistently available, this analysis is intended as a qualitative case study rather than a direct benchmark. Nevertheless, the model assigns distinct affinity predictions to several pairs of compounds with identical connectivity but different stereochemical configurations, suggesting that the proposed representation can capture stereochemistry-dependent variation in drug-target affinity.

Exceptionally, for glutamate [NMDA] receptor subunit 1 (UniProt Q24418), we encounter that (S)-ketamine has been reported to bind with substantially higher affinity than (R)-ketamine, with experimental p*K*_*d*_ values of approximately 6.40 and 5.85, respectively (Jelen et al., 2021). The two enantiomers share identical atomic connectivity and differ only in the configuration of a single stereocenter, yet they exhibit distinct binding behavior at the NMDA receptor.

Table 2 reports the model’s predicted affinities for our manually curated dataset. Specifically, the table lists each protein target together with its corresponding pair of stereoisomeric compound and the predicted affinity assigned to each form. The presence of different predicted values within each pair indicates that the model does not collapse stereoisomers into equivalent representations, but instead produces distinct affinity estimates for molecules that share the same connectivity while differing in stereochemical configuration.

**Table 2:** Predicted affinity differences between stereoisomeric drug pairs across our manually curated set.

| Protein | Isomer A | Isomer B | Affinity (A) | Affinity (B) |
| --- | --- | --- | --- | --- |
| COX-2 | (S)-Ibuprofen | (R)-Ibuprofen | 4.984 | 5.015 |
| SERT | (S)-Citalopram | (R)-Citalopram | 5.182 | 5.046 |
| ADRA1A | (R)-Epinephrine | (S)-Epinephrine | 5.174 | 5.066 |
| β2AR | (R)-Propranolol | (S)-Propranolol | 4.996 | 4.997 |
| NMDAR1 | (R)-Ketamine | (S)-Ketamine | 5.040 | 6.420 |
| GyrB | (R)-Ofloxacin | (S)-Ofloxacin | 5.036 | 5.008 |
| VKORC1 | (R)-Warfarin | (S)-Warfarin | 4.998 | 4.982 |
| ERα | Tamoxifen (Z) | Tamoxifen (E) | 4.999 | 5.001 |
| RAR | Retinoic acid (E) | Retinoic acid (Z) | 5.346 | 5.092 |
| Tubulin | Combretastatin (Z) | Combretastatin (E) | 5.045 | 5.022 |

For the exceptional case, our framework predicts p*K*_*d*_ = 6.41 for (S)-ketamine and p*K*_*d*_ = 5.04 for (R)-ketamine, correctly assigning higher affinity to the (S)-enantiomer. The prediction for (S)-ketamine is in close agreement with the experimental value, while the (R)-form is underestimated by ≈ 0.81 units (see Table 3). Despite this residual deviation in the absolute value of the less active form, the relative ordering of the two enantiomers is preserved and consistent with the experimentally observed stereoselectivity. Since the two compounds are indistinguishable under a purely connectivity-based encoding, this result indicates that the directed-bond message passing scheme is able to leverage stereochemical annotations to differentiate spatial configurations that affect binding.

**Table 3:**
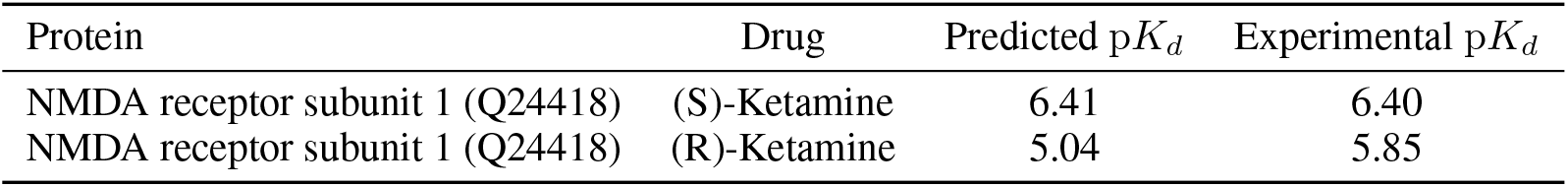
Case study on (S)- and (R)-ketamine at the NMDA receptor (UniProt Q24418). Experimental p*K*_*d*_ values from Jelen et al. (2021).

| Protein | Drug | Predicted $pK_d$ | Experimental $pK_d$ |
| --- | --- | --- | --- |
| NMDA receptor subunit 1 (Q24418) | (S)-Ketamine | 6.41 | 6.40 |
| NMDA receptor subunit 1 (Q24418) | (R)-Ketamine | 5.04 | 5.85 |

## 6 Conclusion

In this work, we presented a stereochemistry-aware framework for drug-target affinity prediction that explicitly encodes three-dimensional configurational information in the molecular representation. By combining a directed-bond message passing neural network on the drug side with frozen ESM-C residue-level embeddings on the protein side, integrated through a cross-attention module, the model achieves state-of-the-art performance on the Davis benchmark while remaining sensitive to stereochemical variation. Furthermore, qualitative evaluation on a newly curated dataset of ten pharmacologically relevant stereoisomeric drug pairs demonstrates the framework’s capacity to discriminate between molecules with identical atomic connectivity. By successfully assigning distinct affinity predictions to different spatial configurations across diverse protein targets, the model proves its sensitivity to stereochemical variations that fundamentally dictate biological activity. These findings support the critical role of stereochemistry-aware representations in developing chemically faithful affinity predictors.

Our evaluation focuses on the Davis dataset, which is restricted to kinase inhibitors and contains a limited number of stereochemically annotated compounds. Extension to broader benchmarks (e.g., KIBA, BindingDB) is needed to assess generalization across target families. The model also relies on one-dimensional protein sequence information; incorporating binding-pocket geometry could further sharpen the discrimination between enantiomers in cases where the magnitude of the affinity gap is currently underestimated.

Stereochemistry-aware affinity prediction can help prioritize the biologically active enantiomer earlier in the drug discovery pipeline, reducing wet-lab effort and avoiding misleading predictions for chiral compounds. As with any in silico predictor, outputs are intended to support, not replace, experimental validation, particularly in safety-critical contexts.

## A Appendix

### A.1 Davis

Table 4 summarizes the presence of stereochemical annotations in the Davis drug set. Among the 68 compounds, 23 contain at least one stereochemical information in their SMILES representation, including chirality (@ and @@) and geometric isomerism (*/* and \). The symbols @ and @@ indicate the local configuration around a chiral center; they encode opposite orientations with respect to the atom ordering in the SMILES string, allowing enantiomeric or diastereomeric forms to be distinguished. In contrast, */* and \ encode directional bonds around double bonds, representing E/Z geometric isomerism caused by restricted rotation. These annotations are therefore important because they preserve three-dimensional molecular differences that can affect drug-target binding affinity.

**Table 4:** Summary of stereochemical information in the Davis drug set. The table reports the number of compounds containing SMILES stereochemical tokens associated with chirality and geometric isomerism.

| Metric | Count |
| --- | --- |
| Total drugs | 68 |
| With @ (chirality) | 14 |
| With @@ (chirality) | 10 |
| With / (E/Z geometry) | 11 |
| With \ (E/Z geometry) | 7 |
| With any stereochemical information (@, /, \) | **23** |

### A.2 Notation

We denote a drug molecule as a directed graph *G* = (𝒱, ℰ) with *N* = |𝒱| atoms and *M* = |E| directed edges (every undirected bond is represented as two directed edges *u* → *v* and *v* → *u*). For each atom *v* we have features 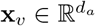 with *d*_*a*_ = 72, and for each directed edge *u* → *v* we have bond features 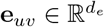 with *d*_*e*_ = 14, both produced by `chemprop’s SimpleMoleculeMolGraphFeaturizer.` We define the *reverse-edge map ρ* such that *ρ*(*u* → *v*) = *v* → *u*.

A protein target is encoded as a sequence of *L*_*p*_ per-residue embeddings produced offline by ESM-C 600m, giving 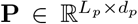 with *d*_*p*_ = 1152. For a mini-batch of size *B*, sequences are right-padded to a common length 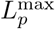 and accompanied by a Boolean mask 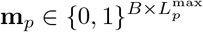 where **m**_*p*_[*b, t*] = 1 iff position *t* of sample *b* is padding.

The model outputs a single scalar *ŷ* ∈ ℝ, the predicted binding affinity, trained against ground-truth *y* with mean squared error.

### A.3 Drug Encoder

The drug encoder follows the directed message-passing scheme of Yang et al. (2019). Hidden states live on edges rather than atoms. Let *W*_in_, *W*_*h*_, *W*_out_ denote the input, hidden, and output linear projections, and *T* the number of message-passing iterations.

#### Input edge embedding

For each directed edge *u*→ *v* we concatenate the source atom features with the bond features and project:

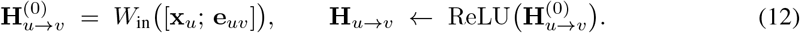

#### Message aggregation

For *t* = 1, …, *T* − 1, the aggregated message into edge *u* → *v* is

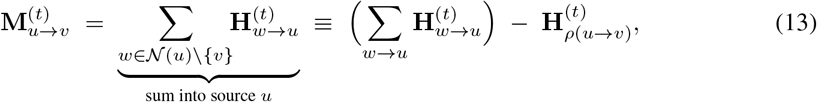

i.e., we sum all messages that arrive at the source atom *u* and subtract the reverse edge’s hidden state to prevent direct backflow along the same bond.

#### Hidden update

At each hidden layer, the representations are updated as per Equation 14:

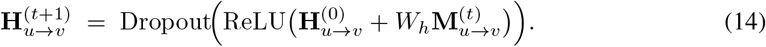

#### Atom readout

After the final iteration, edge messages are scatter-summed onto their target atoms,

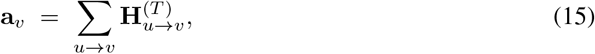

concatenated with the original atom features, and projected:

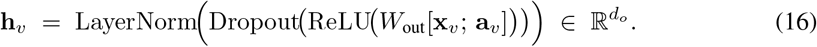

### A.4 Shared Latent Space

Both modalities are mapped into a shared *d*-dimensional latent space via LayerNormalization followed by a small MLP block. Let *L*_*d*_ denote the number of projection layers. The block is

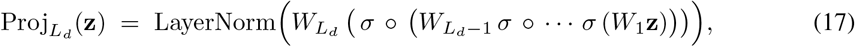

where *σ*(·) denotes ReLU followed by Dropout, the first *L*_*d*_ − 1 linear maps go from input/hidden width to *d*_proj_, and the last linear map goes to the latent width *d*. For *L*_*d*_ = 1 the block reduces to a single LayerNorm(*W* **z**).

#### Drug side

The variable-length per-atom embeddings are first converted to a dense (*B, N* ^max^, *d*_*o*_) tensor with a Boolean atom-padding mask **m**_*d*_. Then

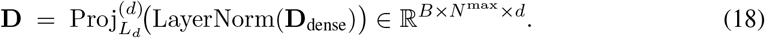

#### Protein side

Similarly, for proteins, we have:

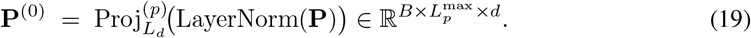

### A.5 Protein Self-Attention

The projected protein representation passes through *L*_ps_ pre-norm Transformer encoder layers:

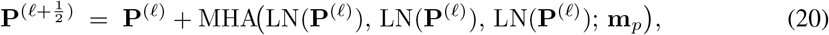

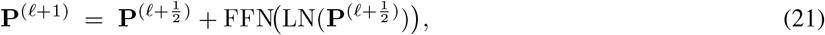

where MHA is multi-head attention with *h* heads, FFN is a two-layer MLP with hidden width 4*d* and a single nonlinearity, and the protein padding mask **m**_*p*_ is used as the key padding mask.

### A.6 Cross-Attention Block

We then feed the projected drug atoms as queries that attend over the contextualized protein residues. Each `CrossAttnBlock` uses *post-norm* residuals (in contrast to the pre-norm protein self-attention above):

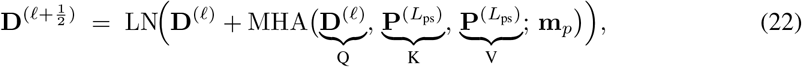

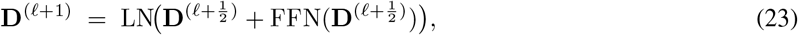

where the FFN is

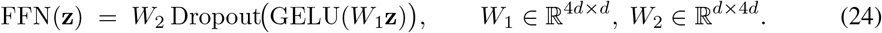

The protein padding mask is supplied as the key padding mask so that padded residues never receive nonzero attention weight from any drug atom. The block is repeated *L*_ca_ times. The drug atom-padding mask **m**_*d*_ is not used inside the block because each query is only ever zeroed out at the readout stage; padded drug rows still carry well-defined intermediate states, but they are discarded by the masked mean below.

### A.7

#### Drug summary

Per-atom contextualized states 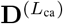 are collapsed to a single fixed-size descriptor by masked mean pooling using the drug atom-padding mask **m**_*d*_:

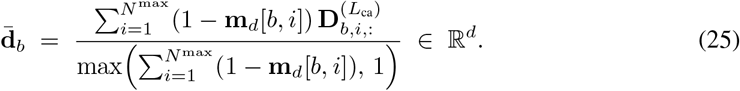

#### Protein summary

The contextualized protein 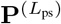 is collapsed by a configurable pooling head:

#### Pooling and Readout

In modes `mean_max` and `mean_max_attn` we compute the masked mean as for the drug side and the masked max as

| Mode | Description | Output dim |
| --- | --- | --- |
| none | no protein descriptor concatenated | 0 |
| mean_max | masked mean and masked max concatenated | $2d$ |
| attn | learnable-query MHA pool | $d$ |
| mean_max_attn | all three concatenated | $3d$ |

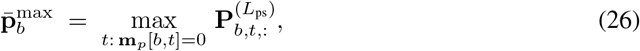

implemented by replacing padded positions with −∞ before torch.max, then defensively replacing any all-padded row with zeros (to handle degenerate inputs).

In modes `attn.` and `mean_max_attn` we additionally use a learnable query token **q** ∈ ℝ^1*×*1*×d*^ (initialized as 𝒩 (0, 0.02^2^), broadcast to batch) and single-step multi-head attention with the protein as both keys and values, followed by a LayerNorm:

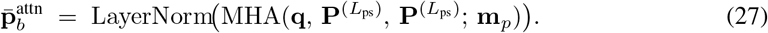

The selected components are concatenated along the feature axis to form 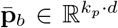 with *k*_*p*_ ∈ {0, 1, 2, 3}.

#### Interaction representation

The final interaction representation is the concatenation of drug and protein summaries:

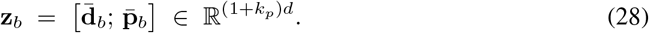

### A.8 Regression Head

A four-layer MLP with fixed widths {512, 256, 64, 1} produces the predicted affinity:

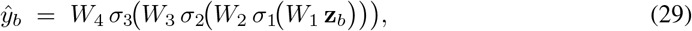

where each *σ*_*i*_ is Dropout ◦ ReLU and the final linear layer has no nonlinearity. The widths are independent of the chosen latent dimension; only the input width (1 + *k*_*p*_) *d* varies.

### A.9 Training Setup

Our model is implemented in PyTorch 2.6 with CUDA 12.1 and trained on a single NVIDIA QUATRO RTX 5600 GPU (24GB). We adopt the Adam optimizer with weight decay and cosine learning rate scheduler and warm-up strategy, and train for up to 200 epochs (around 6 hours of compute time). The complete implementation can be found in [https://anonymous.4open.science/r/DrugTargetAffinityPrediction].

#### Mixed precision

Forward passes execute under `torch.amp.autocast(device_type=“cuda”),` which casts eligible operators (matrix multiplies, attention, convolutions) to FP16 on CUDA devices. The loss is multiplied by a dynamically tuned scalar via `torch.amp.GradScaler` prior to the backward pass to prevent underflow of FP16 gradients; gradients are unscaled before optional clipping and the optimizer step. The scaler is updated each iteration and silently skips steps in which any gradient becomes `NaN` or `Inf`.

